# CoV Genome Tracker: tracing genomic footprints of Covid-19 pandemic

**DOI:** 10.1101/2020.04.10.036343

**Authors:** Saymon Akther, Edgaras Bezrucenkovas, Brian Sulkow, Christopher Panlasigui, Li Li, Weigang Qiu, Lia Di

## Abstract

**Summary:** Genome sequences constitute the primary evidence on the origin and spread of the 2019-2020 Covid-19 pandemic. Rapid comparative analysis of coronavirus SARS-CoV-2 genomes is critical for disease control, outbreak forecasting, and developing clinical interventions. CoV Genome Tracker is a web portal dedicated to trace Covid-19 outbreaks in real time using a haplotype network, an accurate and scalable representation of genomic changes in a rapidly evolving population. We resolve the direction of mutations by using a bat-associated genome as outgroup. At a broader evolutionary time scale, a companion browser provides gene-by-gene and codon-by-codon evolutionary rates to facilitate the search for molecular targets of clinical interventions.

**Availability and Implementation:** CoV Genome Tracker is publicly available at http://cov.genometracker.org and updated weekly with the data downloaded from GISAID (http://gisaid.org). The website is implemented with a custom JavaScript script based on jQuery (https://jquery.com) and D3-force (https://github.com/d3/d3-force).

**Contact:**, City University of New York, Hunter College

**Supplementary Information:** All supporting scripts developed in JavaScript, Python, BASH, and PERL programming languages are available as Open Source at the GitHub repository https://github.com/weigangq/cov-browser.

## Usages & Innovations

Genomic epidemiology comparatively analyzes pathogen genome sequences to uncover the evolutionary origin, trace the global spread, and reveal molecular mechanisms of infectious disease outbreaks including the latest coronavirus pandemic caused by the viral species SARS-CoV-2 (1–4). The unprecedented public-health crisis calls for real-time analysis and dissemination of genomic information on SARS-CoV-2 isolates accumulating rapidly in databases such as GISAID (http://gisaid.org) (5,6). To meet the challenge of real-time comparative analysis of SARS-CoV-2 genomes, we developed the CoV Genome Tracker (http://genometracker.org) with a supporting bioinformatics pipeline. Key features of the CoV Genome Tracker include interactive visualization and exploration of geographic origins, transmission routes, and viral genome changes of Covid-19 outbreaks (Fig 1). A companion comparative genomics website displays the 2003-2004 SARS-CoV and the 2019-2020 SARS-CoV-2 outbreaks in the evolutionary context of their wildlife relatives (1,7).

**Fig 1.**
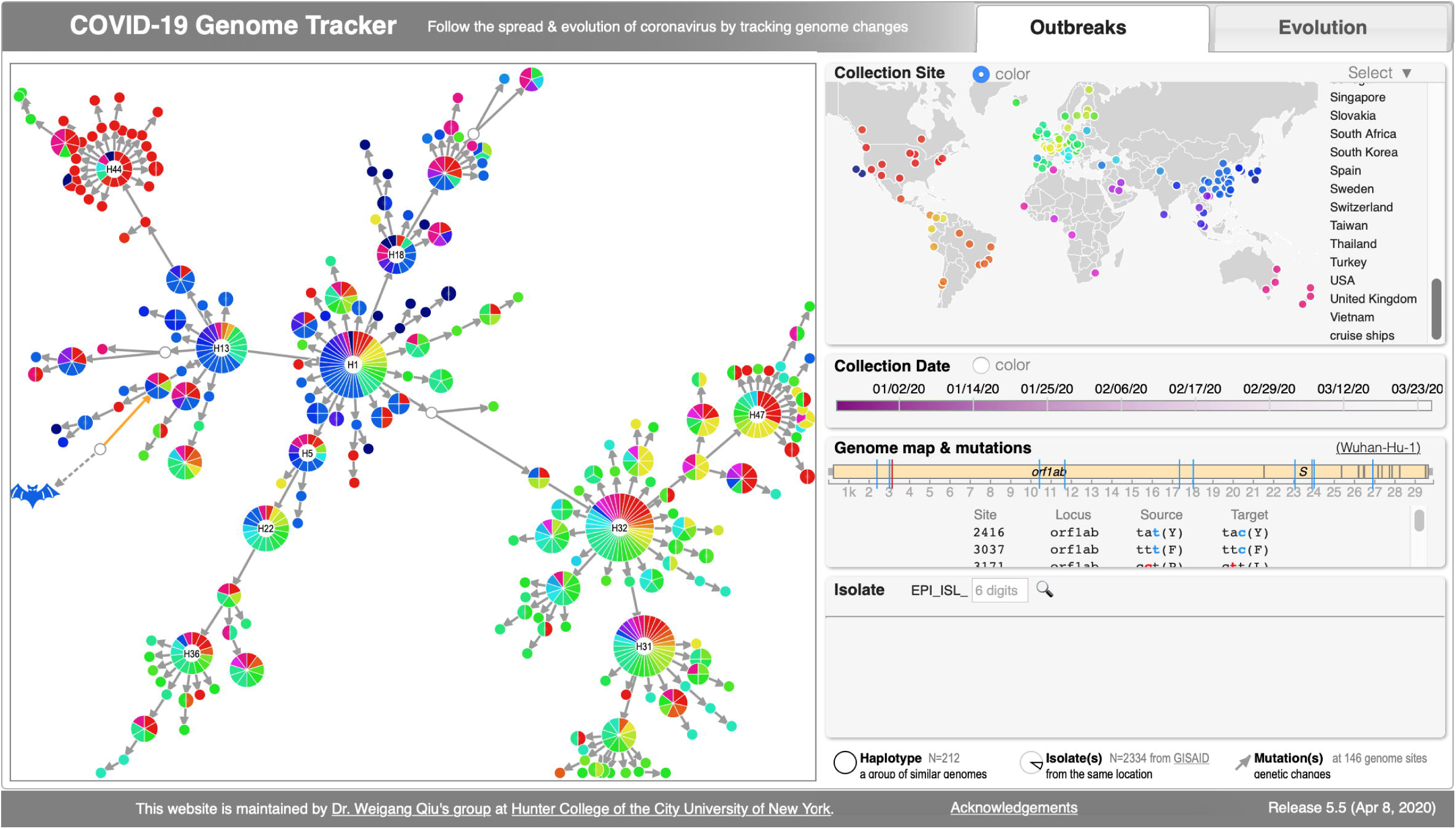
CoV Genome Tracker. uses a maximum-parsimony mutational network (*left panel*) to represent genealogy of SARS-CoV-2 isolates during the 2019-2020 Covid-19 pandemic. The network is interactively linked with geographic origins (color-coded, *top row, right*) and collection dates (*2*^*nd*^ *row, right*) of viral isolates, genomic locations (at n=146 SNP sites) and molecular nature of mutations (*3*^*rd*^ *row, right*), and isolate information searchable by GISAID accession (*4*^*th*^ *row, right*). Colored nodes represent haplotypes (n=212), a unique combination of nucleotides at polymorphic genome positions. Open-circle nodes (n=4) represent hypothetic ancestors. Each slice within a node, occupying one unit area, represents one or more viral isolates (n=2334 genomes downloaded from GISAID as of 3/29/2020) sharing a geographic origin. Thus, node size is an indication of geographic diversity of a haplotype, not the number of isolates. In other words, widely distributed genomes show as large nodes. Large nodes (containing >10 slices) are labeled at the center. Each edge represents one or more mutational changes between a parental and a descendant haplotype. Arrows indicate mutation directions determined according to an outgroup genome (MN996532, strain “RaTG-13”, *bat icon*). The network is consistent with a published one consisting of half the number of genomes (15). However, the maximum parsimony network registers 59 (or 40.7%) sites that have changed more than once (i.e., homoplasy). Causes of homoplasy include sequencing errors, presence of recombination, and the large evolutionary distances between the outgroup and SARS-CoV-2 genomes. Nonetheless, CoV Genome Tracker provides up-to-date genomic changes, helps trace the origin and spread, and facilitates research into virulence mechanisms and clinical interventions on the current and future coronavirus outbreaks.

At the micro-evolutionary time scale, a key distinction of CoV Genome Tracker from the Nextstrain Covid-19 browser (https://nextstrain.org/ncov) (6) is our adoption of a haplotype network – instead of a phylogenetic tree – as the analytic framework as well as the visual guide (Fig 1). A haplotype network offers several advantages over a phylogenic tree. First, at the time scale of days and months, loss and fixation of alleles are rare and the ancestral and descendant genotypes are both present in the population. As such, tree-based phylogenies can be misleading because tree-based phylogenetic algorithms compel all sampled genomes into leaf nodes regardless of ancestral or descendant genotypes, meanwhile introducing hypothetical ancestors as internal nodes. Second, phylogenetic reconstruction typically assumes a mutation-driven process with complete lineage sorting. Violation of these assumptions results in misleading evolutionary relations, for example, when recombination is present or when genes remain polymorphic (8,9). Third, a haplotype network requires less abstract comprehension of evolutionary processes than a phylogenetic tree does. For example, edges of a haplotype network depict genetic changes from a parent to a descendant genome, while branches of a phylogenetic tree represent genetic changes from a hypothetical ancestor to another hypothetical or sampled genome. Fourth, a haplotype network is more scalable than a phylogenetic tree as a visual tool. This is because the total number of nodes of a phylogenetic tree grows linearly with the number of genomes, resulting in a crowded visual space. In contrast, additional genomes add to the size but not the total number of nodes of a haplotype network if they share the same haplotype sequence with previously sampled genomes.

A further innovation of the haplotype network used in the CoV Genome Tracker is the inclusion of an outgroup genome to polarize all mutational changes. Conventional haplotype networks show mutational differences but not mutational directions on edges (10–12). The directed haplotype network in CoV Genome Tracker is thus informative for tracing the origin, following the spread, and forecasting the trend of Covid-19 outbreaks across the globe (Fig 1). To date, one published study and two preprint manuscripts use haplotype networks to represent the genealogy of SARS-CoV-2 isolates (13–15). These networks are however based on a much smaller number of genomes, non-interactive, and non-directional.

At the macro-evolutionary time scale, CoV Genome Tracker provides more in-depth features than the Nextstrain browser on SARS-CoV-2 genome evolution (https://nextstrain.org/groups/blab/sars-like-cov) (6) (Supplemental Fig S1). Modeled after *BorreliaBase* (http://borreliabase.org), a browser of Lyme disease pathogen genomes (16), the comparative genomics browser of CoV Genome Tracker provides analytical features including sequence alignments, gene trees, and codon-specific nucleotide substitution rates. As such, the macro-evolutionary browser facilitates exploring the wildlife origin of SARS-CoV-2, identifying functionally important gene sites based on sequence variability, and understanding mechanisms of genome evolution including mutation, recombination and natural selection (3,4).

## Methods & Implementation

The micro-evolutionary and macro-evolutionary browsers of the CoV Genome Tracker are continuously updated according to the following workflows.

For the Covid-19 genome browser, we download genomic sequences and associated metadata of SARS-CoV-2 isolates from GISAID (5), which are subsequently parsed with a PYTHON script (“parse-metadata.ipynb”; all scripts available in GitHub repository http://cov.genometracker.org). We use a custom BASH script (“align-genome.sh”) to align each genome to an NCBI reference genome (isolate Wuhan-Hu-1, GenBank accession NC_045512) with Nucmer4 (17), identify genome polymorphisms with Samtools and Bcftools (18), and create a haplotype alignment using Bcftools. To minimize sequencing errors, we retain only phylogenetically informative bi-allelic single-nucleotide polymorphism (SNP) sites where the minor-allele nucleotide is present in two or more sampled genomes. To maximize network stability, a custom Perl script (“impute-hap.pl”) is used to trim SNP sites at genome ends where missing bases are common, discard haplotypes with more than 10% missing bases, (optionally) impute missing bases of a haplotype with homologous bases from a closest haplotype (19), and identify unique haplotypes using the BioPerl package Bio::SimpleAlign (20). To root the haplotype network, we include the genome of a closely related bat isolate (RaTG13, GenBank accession MN996532) (1) as the outgroup (using however only nucleotides at the SNP sites present among human isolates).

We use two methods to infer a network genealogy of unique haplotypes. In one approach, we infer a maximum parsimony tree using the DNAPARS program of the PHYLIP package (21). A custom Perl script (“hapnet-pars.pl”) transforms the resulting maximum parsimony tree into a phylogenetic network by replacing internal nodes with the nearest haplotypes where tree distances between the two are zero. Alternatively, we use a custom Perl script (“hapnet-mst.pl”) to reconstruct a minimum-mutation network of unique haplotypes based on the Kruskal’s minimum spanning tree (MST) algorithm implemented in the Perl module Graph (https://metacpan.org/release/Graph). Both Perl scripts polarize the edges of the haplotype network according to the outgroup sequence by performing a depth-first search using the Perl module Graph::Traversal::DFS (https://metacpan.org/pod/Graph::Traversal::DFS). The Perl scripts output a directed graph file in the JavaScript Object Notation (JSON) format. The JSON network file is read by a custom JavaScript, which layouts the website with the JavaScript library jQuery (http://jquery.com) and creates an interactive force-directed rendering of the haplotype network with the JavaScript library D3-force (http://d3js.org).

For the comparative genomics browser of CoV, we download genomes of a human-host SARS-CoV-2 (isolate WIV2, GenBank accession MN996527), a human-host SARS-CoV (isolate GD01, GenBank accession AY278489), and closely related coronavirus isolates from bat hosts from the NCBI Nucleotide Database. We extract coding sequences from each genome and identify orthologous gene families using BLASTp (22). For each gene family, we obtain a codon alignment using MUSCLE and Bioaln (23,24). We reconstruct maximum-likelihood trees for individual genes as well as for the whole genome based on a concatenated alignment of ten genes using FastTree (25). For each gene, we estimate the maximum-parsimony number of nucleotide changes at each codon position using DNACOMP of the PHYLIP package (21). Differences in nucleotide substitution rates between the predominantly synonymous 3^rd^ codon position and the other two codon positions are indicative of forces of natural selection. For example, a higher substitution rate at the 3^rd^ codon position than the rate at the 1^st^ and 2^nd^ positions indicates purifying selection while a higher or similar rate at the 1^st^ and 2^nd^ codon positions relative to the rate at the 3^rd^ codon position suggests adaptive diversification (e.g., at the Spike protein-encoding locus) (2). The CoV comparative genomics browser is developed with the same software infrastructure supporting *BorreliaBase* (http://borreliabase.org), a comparative genomics browser of Lyme disease pathogens (16).

## Conclusion & Future Directions

In summary, the CoV Genome Tracker facilitates up-to-date and interactive analysis of viral genomic changes during current and future coronavirus outbreaks. The CoV Genome Tracker uses a haplotype network, a more accurate and scalable model than a phylogenetic tree to analyze and visualize genomic changes in the rapidly evolving SARS-CoV-2 population (6). We improved upon conventional haplotype networks by resolving the direction of mutational changes based on an outgroup genome (10,12). Future development will include implementing probabilistic network algorithms such as maximum parsimony probability (10,11), developing methods for testing network accuracy and stability, analyzing association between genomic changes and network characteristics (e.g., association between the number of nonsynonymous mutations and the in- and out-degrees of nodes), performance optimization, usability improvements, and incorporating a mechanism for community feedback.

## Declaration

### Availability of website & source codes

CoV Genome Tracker is publically available at http://cov.genometracker.org. All source codes are released as Open Source and available at https://github.com/weigangq/cov-browser. The repository contains BASH, Perl, Python, R, and JavaScript codes for data processing pipeline, network reconstruction, and web development.

### Authors’ Contributions

S.A. implemented the genome processing pipeline and drafted the manuscript. E.B. developed the workflow for downloading and parsing data from the GISAID database. B.S. performed network stability analysis and contributed to website design. C.P. prepared and maintains online documentation. L.L. contributed to network analysis, drafting manuscript, and online documentation. W.Q. conceived the project, developed and implemented the network algorithm, and drafted the manuscript. L.D. developed the meta-data pipeline, designed the website, implemented JavaScript codes, and prepared the figures.

## Acknowledgements

We gratefully acknowledge the authors, originating and submitting laboratories of the sequences from GISAID’s EpiCoV™ Database on which this research is based. All submitters of data may be contacted directly via www.gisaid.org. We thank Desiree Pante, Bing Wu, and Ramandeep Singh for participation in data entry. We thank Dr Yozen Hernandez for system administration of computer networks. We thank Jonathan Sulkow for contributing to webpage design.

