## Supplemental Figure S1 for "CoV Genome Tracker: tracing genomic footprints of Covid-19 pandemic"

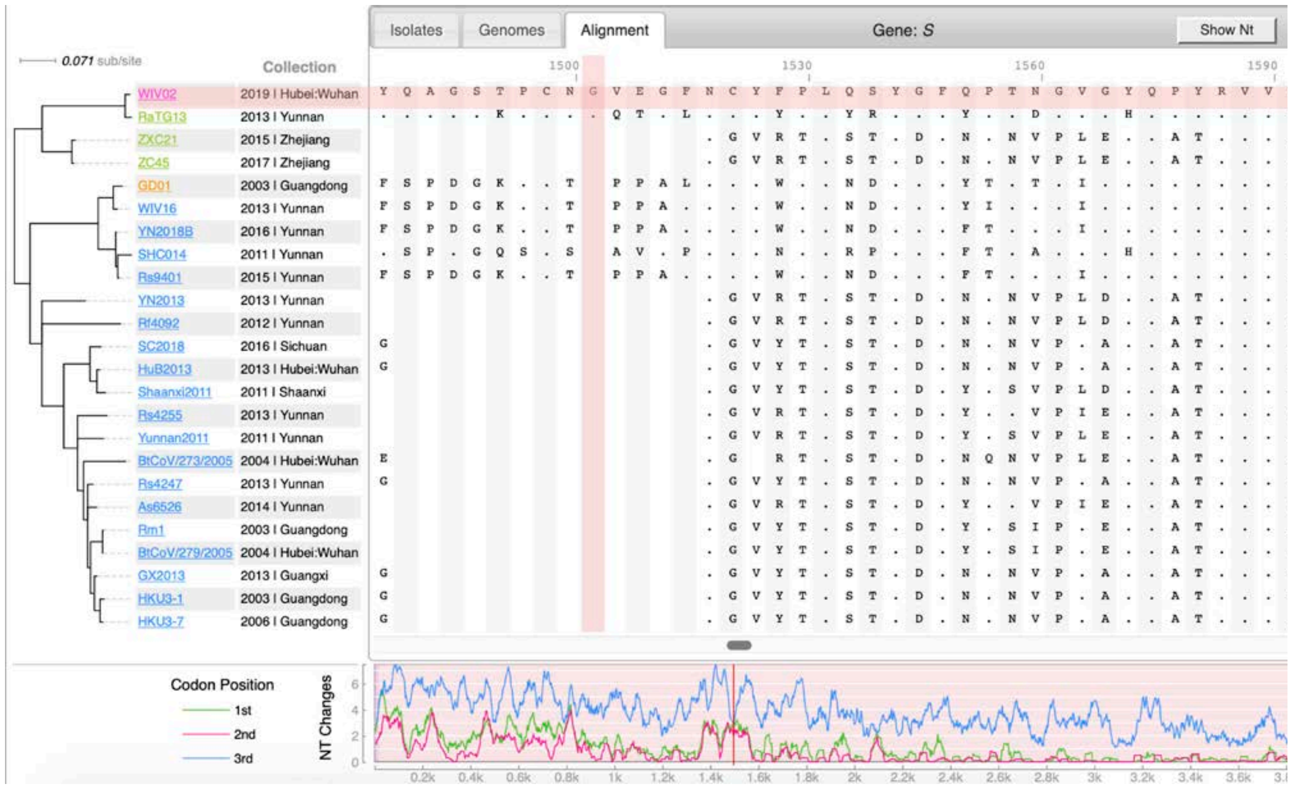


**Fig S1. Macro-evolutionary browser of coronavirus genomes.** This is a companion genome browser of SARS-CoV (SARS) & SARS-CoV-2 (COVID-19) genomes in the evolutionary context of their respective animal-host relatives in China. Here, aligned amino-acid sequences (*right panel*) of the Spike protein – showing part of the highly variable receptor binding domain (RBD) – are arranged according to the sequence phylogeny (*left panel*). All sequences are compared to the Covid-19 strain (WIV02) at the top, with "." representing the same residue, letters representing a difference, and blanks representing alignment gaps. At the bottom, we show nucleotide substitution rates at three codon positions, as a way to show gene sites under positive selection (e.g., 5'-end, where rates at 1$^{st}$ and 2$^{nd}$ codon positions reach to the level at 3$^{rd}$ codon position), as well as those under negative selection (3'-end, where changes are mostly on the 3$^{rd}$ synonymous codon positions). This macro-evolutionary browser is designed after *Borreliabase*, a browser of Lyme disease pathogen genomes we developed and maintain (Di et al. 2014). This browser helps, e.g., the design of vaccines against a specific CoV strain, a group of CoV strains, or all future CoVs spilling over from animal reservoirs.
